# Actin turnover required for adhesion-independent bleb migration

**DOI:** 10.1101/2022.04.01.486735

**Authors:** Calina Copos, Wanda Strychalski

## Abstract

Cell migration is critical for many vital processes, such as wound healing, as well as harmful processes, like cancer metastasis. Experiments have highlighted the diversity in migration strategies employed by cells in physiologically relevant environments. In 3D fibrous matrices and confinement between two surfaces, some cells migrate using round membrane protrusions, called blebs. In bleb-based migration, the role of substrate adhesion is thought to be minimal, and it remains unclear if a cell can migrate without any adhesion complexes. We present a 2D computational fluid-structure model of a cell using cycles of bleb expansion and retraction in a channel with several geometries. The cell model consists of a plasma membrane, an underlying actin cortex, and viscous cytoplasm. Cellular structures are immersed in viscous fluid which permeates them, and the fluid equations are solved using the method of regularized Stokeslets. Simulations show that the cell cannot effectively migrate when the actin cortex is modeled as a purely elastic material. We find that cells do migrate in rigid channels if actin turnover is included with a viscoelastic description for the cortex. Our study highlights the non-trivial relationship between cell rheology and its external environment during migration with cytoplasmic streaming.

## 1. Introduction

Single cell migration is an almost ubiquitous phenomenon in eukaryotic biology that serves many important physiological roles including embryonic development, immune response, and wound healing [1]. Cells use a variety of biophysical mechanisms to migrate that vary depending on their external environment [2]. For example, mesenchymal migration employed by cells such as keratocytes and fibroblasts on a flat 2D substrate is characterized by actin polymerization at the leading edge and substrate adhesion [3–5]. In contrast, some cells in 3D use an amoeboid mode of motility, where cells have a round morphology and lack mature adhesions and actin stress fibers [6,7]. Amoeboid migration plays important roles in developmental biology and immune system function [8,9]. Additionally, certain tumor cells can transition between mesenchymal and amoeboid migration modes and thereby increase cancer invasiveness [1]. Some cells have even been shown to switch migration modes after biochemical or mechanical stimulation [10–12].

The focus of this manuscript is on a relatively novel migratory mechanism of rounded cells that do not rely on cell-surface adhesion for efficient migration in complex 3D environments [13,14]. We refer to this form of locomotion as *adhesion-independent amoeboid movement*. This migration mode is relevant for the migration of leukocytes through the endothelial barrier out of blood circulation to the location of damaged tissue during wound healing [8,15], and the *in vivo* migration of various cells in developing embryos [16], metastatic cancer [1,17], and immune cells migrating through tissue while patrolling for pathogens [18].

Amoeboid cells move much faster and are more autonomous from their extracellular environment than mesenchymal cells in the sense that they adapt to their environment instead of remodeling it [13]. Locomotion of amoeboid cells does not depend on adhesive ligands, and they are able to migrate efficiently in a non-frictional manner even in suspensions or artificial materials [15,19]. In the adhesion-independent amoeboid mechanism, cells find the path of least resistance through the extracellular matrix by picking larger pores in the matrix over small ones [20], actively deforming their cell body and/or transiently dilating the pore in order to pass it through [21–23]. Charras and Paluch hypothesized that a cell exerts forces perpendicularly to the substrate such that it can squeeze itself forward using membrane protrusions called *blebs* [24]. This phenomenon has been termed ‘chimneying’ [25] (authors observed cells migrating between two glass coverslips), in reference to a technique used by mountain climbers. Exactly how cellular forces are transmitted to the substrate in order to produce migration remains an open question.

In this work, we consider amoeboid cell migration in the case where the leading edge protrusion is generated by a bleb, a spherical membrane protrusion characterized by a delamination of the actomyosin cortex from the cell plasma membrane [26]. When the actin cortex is separated from the membrane, tension from actomyosin contractility is no longer transmitted to the membrane in the delamination region, and pressure is locally reduced [27]. Cytoplasmic content then streams from the cell body into this region, expanding the round membrane protrusion. Bleb initation can occur by either a localized loss in membrane-cortex adhesion proteins or local-rupture of the cortex [24,28]. Actin, myosin, and associated proteins eventually reform under the naked membrane to drive bleb retraction and complete the life-cycle of a bleb. Blebbing cell migration has been observed in a number of cell types, such as amoebas, zebrafish germ layer progenitor cells, and Walker 256 carcinosarcoma cells [16,29,30].

Several computational models have been developed to investigate the relative importance of key factors in confined migration, such as geometry of the environment, actomyosin contractility, role of nucleus, and type of leading edge protrusion [31–33]. The aim of such computational modeling is to complement the experimental findings and propose mechanisms for generating internal forces and transmitting these forces to the surface in order to produce traction. A detailed hybrid agent-based/finite-element model of cancer cell motility was presented in [32]. The authors consider a number of migratory mechanisms including pressure-driven (amoeboid) and actin-rich (mesenchymal) protrusions on flat surfaces, channels, and discontinuous 3D-like environments with varying levels of cell-surface adhesion. The authors found that in the absence of any cell-surface adhesions only cells exhibiting the amoeboid mechanism could migrate efficiently in a discontinuous environment. A similar type of model was proposed in [33] to quantify conditions for motility modes for a cell migrating through an elastic extracellular matrix. One limitation of both models is that intracellular fluid flow is not incorporated (i.e., intracellular pressure is treated as constant). The model in [31] does include intra-and extracellular fluid mechanics, but results focus on the role of nuclear stiffness during amoeboid cell migration. The theory of active gels has also been used to model confined cell migration [34,35]. For example in [35], the authors show that motion can occur in confinement when the cell cytoplasm is modeled as a polymerizing viscoelastic material.

A natural question to ask is whether a confined environment is even necessary for cell migration in a fluid environment. Several groups have developed models to determine conditions when a cell can “swim” in low Reynolds number fluid. The Scallop Theorem states that time-symmetric motion cannot achieve net displacement in Stokes flow [36]. In order to swim in fluid, many prokaryotes (and sperm, paramecia and some eukaryotes) use a beating flagella or cilia to migrate. Such flagellated organisms achieve self-propulsion through periodic flagellar bending waves [37,38]. Experiments have shown that amoebae and neutrophils are also able to swim [39], and several models of amoeboid cell swimming have been developed (reviewed in [40]). A model for bleb-based swimming modeled the cell as two spheres submerged in fluid that can expand or contract radially that are connected by an extensible arm in [41]. An amoeboid cell representative of *Dictyostelium discoideum* immersed in fluid was shown to swim through shape changes and membrane tension gradients in [42,43].

In [44], the authors developed a model similar to ours in that it includes a blebbing cell immersed in Stokes fluid with the model equations solved for using the method of regularized Stokeslets [45]. The authors showed that in their model the cell was able to swim because membrane deformations during bleb expansion differed from those during bleb retraction. Results showed that migration speed was optimal at an intermediate confinement level. This model was then used to predict that the optimal gap size increases with weakening adhesion between the cell membrane and actin cortex, which was experimentally verified in [46].

Here, we present a dynamic computational model of adhesion-independent cell migration using cycles of bleb expansion and retraction. Our model is formulated using the method of regularized Stokeslets [45] to handle the fluid-structure interaction. Our model differs from previous work in that we consider intra and extra-cellular fluid flows, the actomyosin cortex is modeled as a poro-viscoelastic material, and various channel geometries are considered. Our results show that cyclic pressure gradients from blebbing together with cortical actin dynamics result in cell shape changes such as expansion and contraction of the cell body. However, the shape change pattern is nearly reversible and does not result in sustained net locomotion, even in confined environments. In exploring design principles for locomotion, the channel width is varied and an asymmetrical wall geometry is considered. Neither one of these endeavours improved cell movement. However, introducing actin turnover did produce sustained net locomotion in both suspensions and confined environments. Our results show that confinement enhances locomotion speed for most channel geometries. Speficially, if the wall has large crevices, a cell can become stuck and bleb vertically within a channel gap. Simulation results also show that migration speed increases with actin turnover.

The paper is organized as follows. In Section 2, we describe the model of the cell, channel, and bleb life cycle. We describe the computational algorithm to solve and implement the model equations. In Section 3, the model is first simulated using a poroelastic cortex using different bleb sizes as well as channel geometries. Next, we consider blebbing with cortical actin turnover as modeled by the Maxwell viscoelastic constitutive law in unconfined and various confined geometries. The effect of actin turnover on cell migration speed is also quantified. A discussion of results and conclusion remarks are provided in Section 4.

## 2. Materials and Methods

We build a computational model of a cell placed in a microfluidic rigid channel undergoing cycles of bleb expansion and contraction driven by intracellular fluid flows as illustrated in Fig. 1. The model is two-dimensional in that it captures the motion of the cell in the horizontal direction of motion as well as the channel height. We assume that the flow in the third dimension across the channel is negligible. The motion in the horizontal direction is due to extracellular fluid flows induced by cell shape changes. Our model has three sub-cellular components: an elastic plasma membrane, a contractile actomyosin cortex, and the cell cytoplasm. The cell cytoplasm is assumed to be a viscous fluid enclosed in the thin actin cortex and plasma membrane. The cortex is modeled as a thin 1D porous (visco)elastic material and its position is denoted by ***X***^cortex^(*s, t*) where *t* is time and *s* is the local parametric coordinate on the structure. The plasma membrane is described as an incompressible 1D elastic outer layer with position ***X***^mem^(*s, t*). The cortex is bound to the plasma membrane via membrane-anchoring proteins modeled here as elastic links [47]. When a bleb is initiated, membrane-cortex attachment links are removed and the cell front expands due to the emergent fluid pressure gradient. As the cortex reforms at the bleb site, membrane-cortex attachment links reform and the cell front retracts. Since the focus is on adhesion-independent migration, there are no physical links between the cell and the extracellular environment.

**Figure 1.**
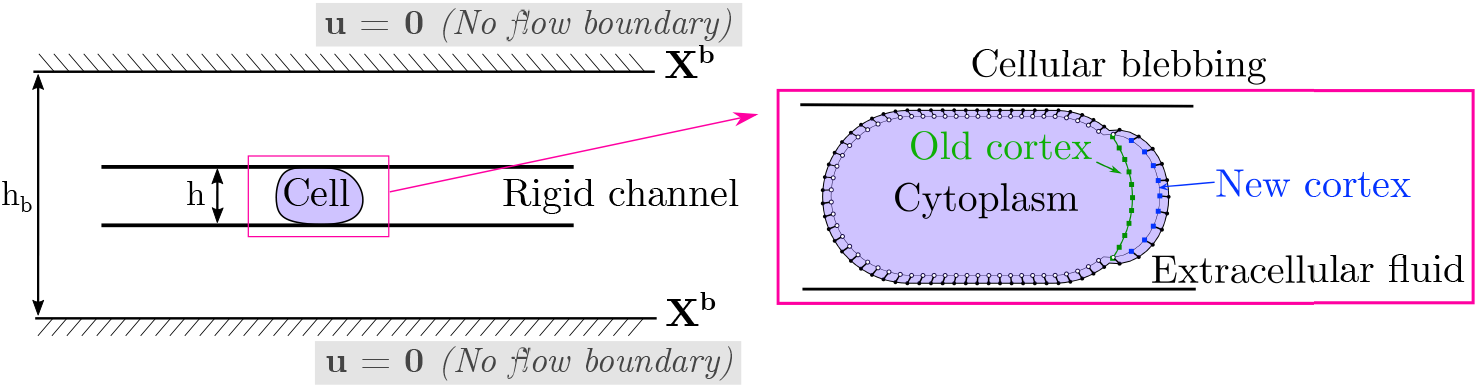
Schematic of blebbing cell in a channel. In our model, a cell is compressed within a rigid channel of adjustable wall geometry and height. No flow boundary conditions are enforced at an outer boundary ***X***^b^. The default setup is a confinement height of *h* = 13 *μ*m for a 20 *μ*m diameter cell with an outer boundary of height *h*_b_ = 32 *μ*m. The cell model consists of an incompressible plasma membrane (black dots), a (visco)elastic actomyosin cortex (white, green, and blue square points), and a viscous cytoplasm. The lines connecting membrane and cortex points represent membrane-cortex attachment links.

### 2.1. Equations of Motion

Movement in viscous fluid at zero Reynolds number is governed by Stokes equations due to the small length scales at the cellular level [36]. In our model, the external forces applied to the fluid are due to the deformations of the plasma membrane, the membrane-cortex attachment links, the viscous drag with cortex, as well as a repulsive steric interaction with the top and bottom channel walls:

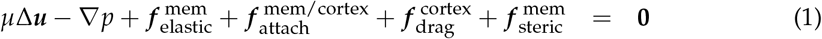

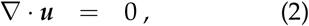

where ***u*** is the fluid velocity, *p* is the fluid pressure, and *μ* is the fluid viscosity. Expressions for these cellular forces are provided below. We use the convention of lower case letters indicating fluid quantities while upper case letters indicate forces and positions of structures.

#### Outer plasma membrane and actomyosin cortex

We consider two rheological descriptions for the membrane and cortex contours. Each contour experiences forces due to either elasticity or viscoelasticity. Let 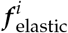 denotes the elastic force density on the membrane and cortex

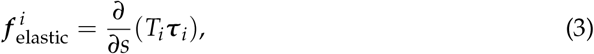

where ***τ**_i_* denotes the unit tangent vector to the closed curve Γ^*i*^ = ***X**^i^*(*s, t*) = ***X***^mem^(*s, t*) or ***X***^cortex^(*s, t*). The tension *T* is given by

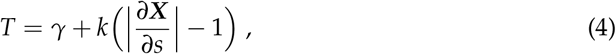

which describes a linearly elastic spring with stiffness *k* and resting tension *γ*. The plasma membrane and actomyosin cortex have their own characteristic stiffness and resting tension (see Table 1). When the membrane and cortex are modeled as viscoelastic structures, we use a Maxwell model to capture stress relaxation of the actomyosin cortex due to actin filament rearrangement within the cortex [48]. Following the approach in [49], a viscoelastic structure is modeled as a purely elastic spring (see Eqs.(3)–(4)) whose reference configuration ***X***_0_ relaxes to the current configuration ***X*** over time with the derived expression

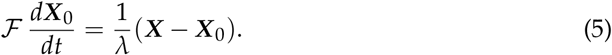

**Table 1.**
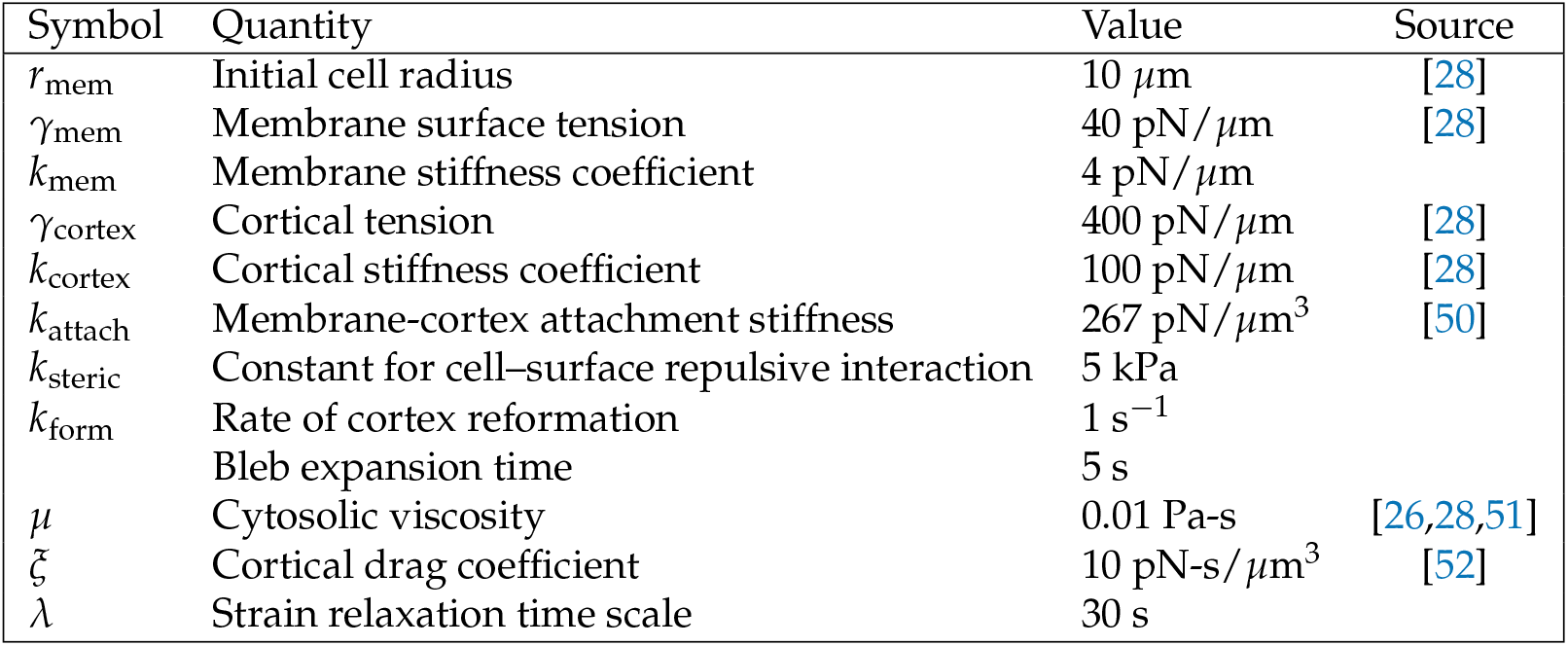
Model parameters.

Here, 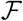 is the deformation gradient tensor 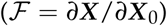 and *λ* is the strain relaxation timescale. In the limit of small strain, the authors in [49] show that the update equation for the reference configuration in Eq. (5) together with the elastic force in Eq. (3) agrees with the Maxwell model for viscoelasticity.

#### Attachment between the membrane and cortex

Membrane-cortex attachments are modeled as elastic springs that connect the plasma membrane to the underlying actin cortex with a force density given by

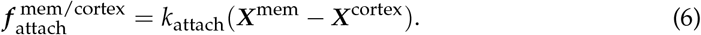

#### Viscous cell cortex drag

The drag force on the cell cortex is balanced by (visco)-elastic forces within the cortex and elastic forces from cortex attachment to the plasma membrane:

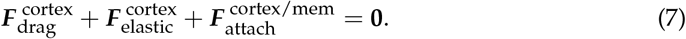

The cortical drag is defined as 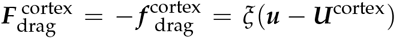 where *ξ* denotes the viscous drag coefficient and 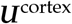 is the cortex velocity.

#### Cell-surface interaction

The cell interacts with the channel walls through a repulsive force due to contact with the surface:

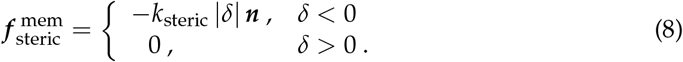

Note that the steric force is only nonzero when the membrane location exceeds the top or bottom channel walls. Here, *δ* is the vertical distance from the plasma membrane to the channel walls, ***n*** is a unit vector in the outward normal direction, and *k*_steric_ is the stiffness of the steric interaction.

#### Formation and retraction of a cellular bleb

In order to account for reformation of the actin cortex within the bleb, we include an additional numerical contour to represent the new cortex in the bleb. Fig. 1(b) shows the location of the old and new cortex points and the location of membrane-cortex attachment links during blebbing. Cortical elasticity is multiplied by local density *ρ* on the new and old cortex. A bleb is initiated by setting the density of the new cortex to zero in a small region at the front of the cell. Adhesive links between the old cortex and the membrane are removed but are maintained in the new cortex. Since the elasticity is zero when a bleb is initialized, the new cortex points stay close to the membrane in the growing bleb. We specify a time for bleb expansion of 5 seconds to allow a large bleb to form at the cell front. After 5 seconds, a bleb moves to the retraction phase.

During bleb retraction, the density of the new (*ρ*_new_) and old cortex (*ρ*_old_) are updated over time according to the following equations:

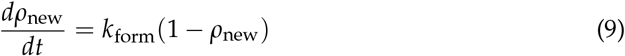

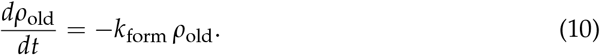

The rate of cortex reformation at the bleb site is *k*_form_. Note that densities of the new and old cortex are assumed to be spatially uniform in their respective locations. In the bleb retraction phase, *ρ*_new_ increases over time while *ρ*_old_ monotonically decreases. Once the density on the old cortex reaches a critical value of 0.2 during bleb expansion, the bleb switches from retraction to expansion by resetting the density on the old cortex to a value of 1 and the density of the new cortex to 0.

Once a bleb cycle is completed (i.e., after the density on the old cortex reaches a value of 0.2), the old cortex is reset to coincide with membrane points at the start of the new cycle of bleb expansion. The process ensures a re-calibration at the beginning of a new cycle.

#### Update equations

Given a configuration of the membrane and cortex structures, forces at every location on the structures are computed as described above, and then the pressure and velocity of the fluid, along with velocity of the membrane and cortex are obtained by solving Eqs. (1)–(2) and Eq. (7). The position of each structure is updated according to their own respective velocities:

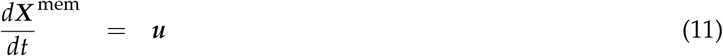

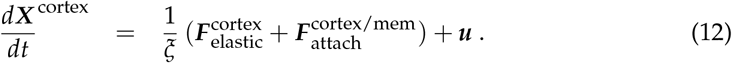

The viscoelastic response of the membrane and cortex structures require each an additional equation for the stress relaxation of the structures, namely Eq. (5).

### 2.2. Numerical Method

Given an initial configuration of the plasma membrane and cortex, the one-dimensional contours are discretized into a finite number of nodes. At every node on the membrane and cortex, forces are computed according to constitutive laws provided in the previous section. After forces are numerically computed, we use the method of regularized Stokeslets [45] to solve for the fluid velocity and pressure in Eqs. (1)–(2). In free space, the fluid velocity at the membrane and cortex structures is

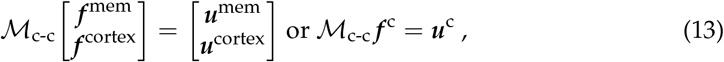

where

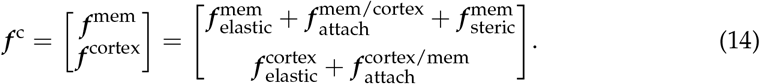

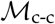 denotes the regularized Stokeslet matrix with entries

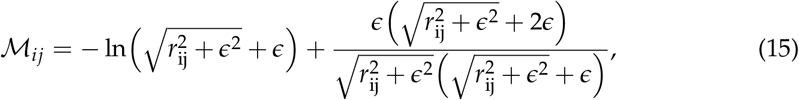

where *r*_ij_ = |***X**_i_* – ***X**_j_* | and the regularization parameter *ε* = 1.5Δ*s* which maps smeared cellular forces to fluid velocities at the cellular structures. Here, we use the 2D blob function *ϕ_ε_* (***x***) from [45] to spread or regularize a point force density over a small ball around a point ***x***. Values for numerical parameters such as Δ*s*, the grid spacing of the discretized membrane contour, are listed in Table 2. Once the fluid velocity is known, the position of the immersed cellular structures are update using the forward Euler time integrator scheme applied to Eqs. (11)–(12).

**Table 2.** Computational and discretization parameters.

| Symbol | Quantity | Value |
| --- | --- | --- |
| $N_{ib}$ | Membrane/cortex mesh size | 134 |
| $\Delta s$ | Initial structure grid step size | $2\pi r_{\text{mem}}/N_{ib}$ |
| $\Delta t$ | Time step size | 1e-4 s |
| $\epsilon$ | Regularization parameter | $3\Delta s/2$ |

Although the force balance in Eqs. (1)–(2) ensure zero sum of forces, the introduction of cell-surface interaction can lead to a force imbalance. In 3D, the first term in the Stokeslet decays like 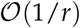, where *r* is the distance from a point force, whereas in 2D, this term decays like 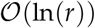. Therefore, even a numerically negligible error in sum of forces can result in ||***u*** || → ∞ as *r* → ∞. To ensure the forces sum to zero, we enforce a no flow boundary condition far away from the physical domain (see Fig. 1 (right)). Although we could have imposed the boundary condition directly on the channel walls as in [44], our approach allows us to avoid resolving thin fluid boundary layers between the cell and channel wall from satisfying a no-slip boundary condition. Our approach satisfies a no-penetration boundary condition ***u*** · ***n*** = 0 on the channel wall and was previously used in [49] to simulate cell deformation in a microfluidic channel with an imposed background flow. To obtain the velocity on membrane and cortex nodes, the linear system in Eq. (13) must then be modified to

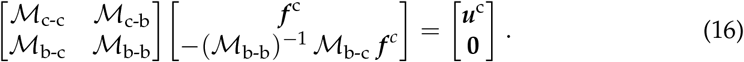

The notation 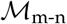 denotes the regularized Stokeslet velocity matrix mapping smeared forces at locations ***X***^n^ to velocities at locations ***X***^m^. For example, 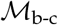 describes the effect of cellular forces at ***X***^c^ to evaluate velocities of the outer boundary channel ***X***^b^. Note that the method in Eq. (16) ensures that there is no fluid flow at the boundary location, i.e. ***u***^b^ = **0**. Alternatively, one can rewrite Eq. (16) for the fluid flow at cellular nodes ***X***^c^ as

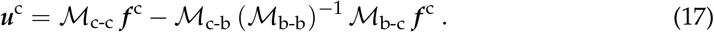

Pressure is computed as follows,

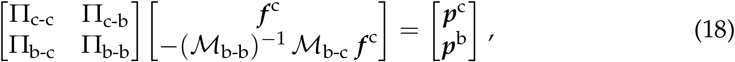

where Π_m-n_ represents the regularized Stokeslet pressure matrix which maps regularized forces at locations ***X***^n^ to pressure at locations ***X***^m^ [45]. Thus, pressure along the cellular locations ***X***^c^ is given by:

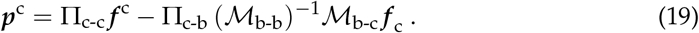

To compute the pressure at arbitrary locations ***X***^q^, which include the cell and external boundary wall forces,

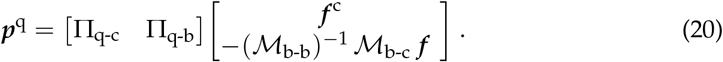

#### Re-meshing algorithm

In the limit of small relaxation, our model for viscoelasticity describes a fluid rather than solid; the method does not guarantee to preserve the mesh spacing as the material deforms. Thus, for large deformations, in the case of moving, deforming structure, a re-meshing algorithm maintains resolution of the discretized structures. Here, the protocol is to re-mesh when a bleb cycle is completed (i.e., the density on the old cortex reaches a value of 0.2). In order to re-space the nodes on the cortex and membrane structures uniformly and preserve their strain, a periodic spline function is used to construct differentiable functions from the position of the discrete nodes. The integral of these differentiable functions yields the arclength of the closed curve, 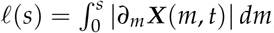, as a function of current Lagrangian coordinate for the deformed configuration. The inverse map from the Lagrangian coordinate to the corresponding arclength is computed using another periodic spline function, *s*(*ℓ*). Lastly, we define a new equally-spaced arclength function and compute the new parameteric coordinate on the arclength by evaluating the previously formed function. Similarly, a periodic spline function is formed for the tension, *γ*(*s*), and it is evaluated at the new parametric coordinate locations, *γ*(*s*^new^). A similar re-meshing algorithm was implemented and tested in [31].

## 3. Results

First, we simulate bleb expansion and retraction when the immersed cell is unconfined (immersed in viscous fluid). A bleb is initiated by specifying a region on the right side of the cell where the density of new cortex is set to zero, and adhesive links between the old cortex and membrane are removed. This region is defined as the bleb neck. Forces from the old cortex (due to elasticity) are not transmitted to the membrane in the bleb neck, and the corresponding forces from the new cortex are zero during bleb expansion (as described in Section 2.1). This leads to a localized pressure gradient and fluid flow that expands the cell membrane and forms the protrusion.

Membrane position at several time values during one cycle of bleb expansion and retraction are shown in Fig. 2 with a bleb neck size of 16 *μ*m. Because the motion of the membrane appears to be approximately reciprocal, we do not expect significant migration (or swimming) and simulation results show almost no displacement of the cell after one bleb cycle. Since significant motion in confinement was reported using a similar model of blebbing [46], we explore the possibility of confined migration with our model. We return to the case of a cell freely swimming using cycles of blebbing in Section 3.2.

**Figure 2.**
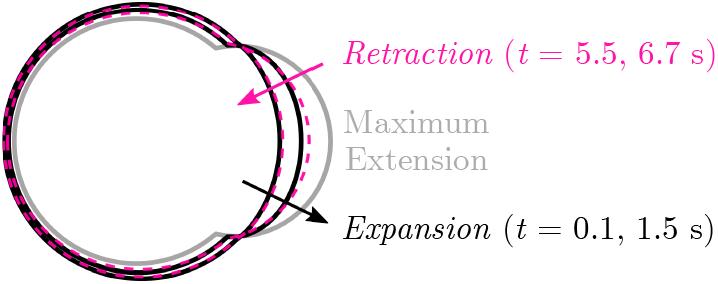
(Nearly) Reciprocal motion in one bleb cycle. The plasma membrane position during one bleb cycle for a cell with an elastic membrane and cortex (no cortical turnover). Membrane position is labeled in black during bleb expansion and by a dashed magenta line during bleb retraction. The bleb is fully expanded at *t* = 5 s.

### 3.1. Elastic Actomyosin Cortex Insufficient for Sustained Locomotion in Confinement

Next, we simulate cycles of blebbing when the cell is placed within a rigid channel. The size of the bleb and the environment’s physical properties are varied and the resulting deformations and motion are shown in Fig. 3 and Fig. 4. Surprisingly, we find that neither increasing the bleb size nor introducing geometrical asymmetries in the channel can produce persistent forward locomotion.

**Figure 3.**
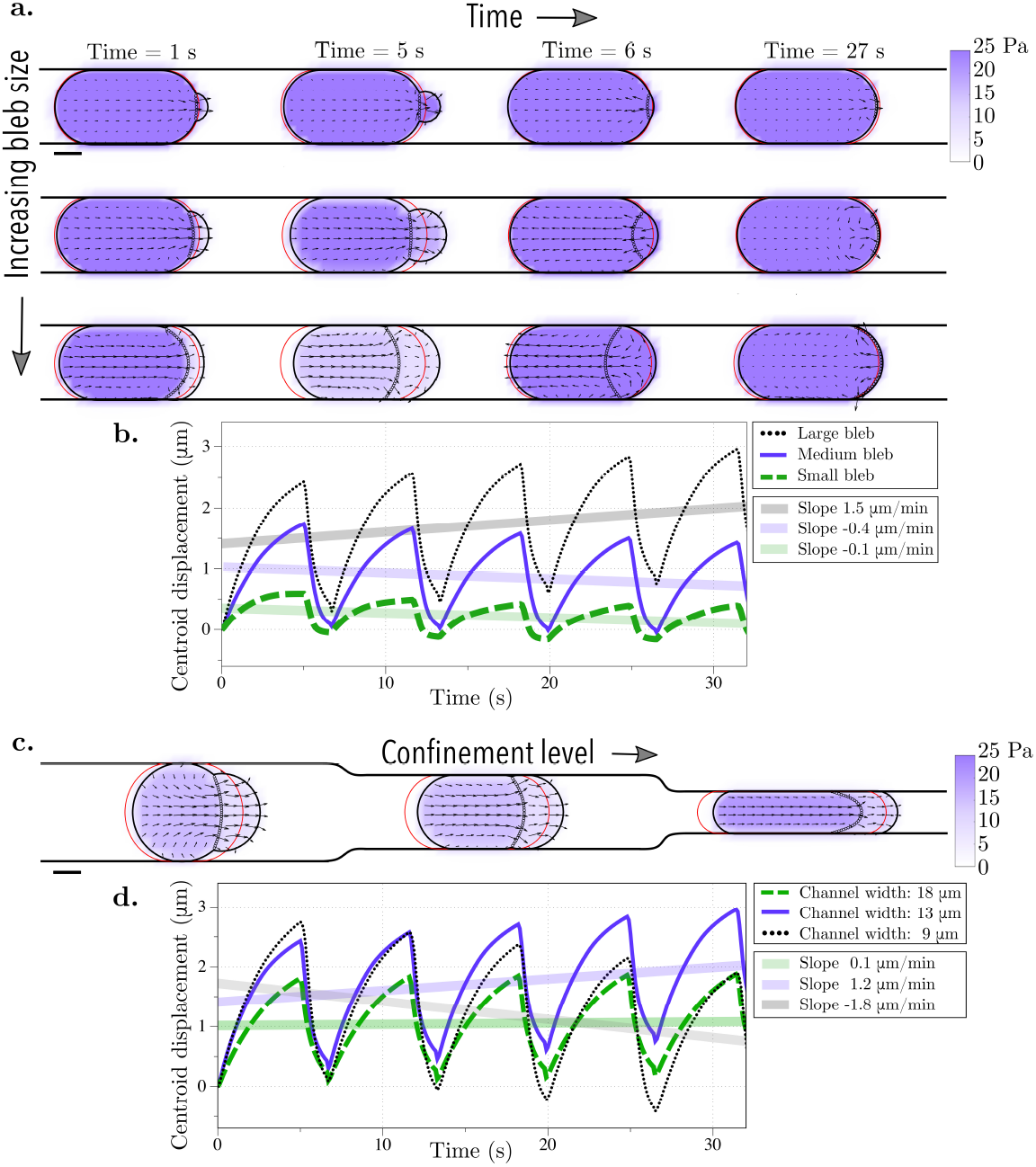
Bleb cycle in a straight rigid channel. Bleb expansion and retraction over one cycle (approximately 7 s) and after 4 bleb cycles (27 s) for (a) three different bleb sizes (bleb neck sizes of 4, 8, and 16 *μ*m) and (c) three different channel widths (18, 13, and 9 *μ*m). The vector field represents the intracellular fluid velocity, and the scalar color field represents cytosolic pressure. The initial position of the membrane is shown in red. The current position of the membrane is shown in black, and the position of the old cortex are denoted with black circles. (b) & (d) The straight line in the centroid horizontal displacement is a least squares fit to the data over 4 bleb cycles. (b) The slopes for the small bleb, medium, and large bleb, respectively, are −0.1, −0.4, and 1.5 *μ*m/min. (d) The slopes for linear fits of simulation data for channel heights of 18, 13, and 9 *μ*m are 0.1, 1.2, and −1.8 *μ*m/s, respectively.

**Figure 4.**
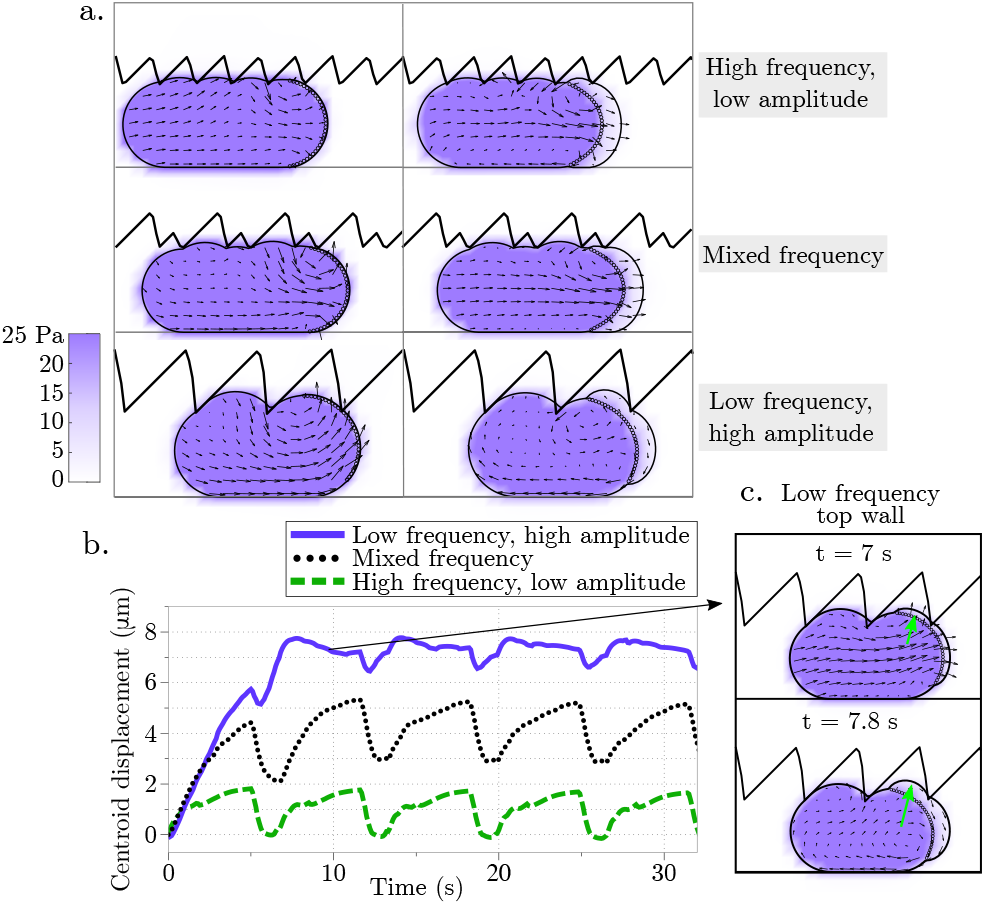
Displacement during blebbing in an asymmetrical rigid channel. (a) Cell position as the bleb expands at two time points (left panels at 19.9 s and right panels at 20.9 s) for different geometries of the top channel wall: high frequency/low amplitude, mixed frequency, and low frequency/high amplitude sawtooth. The vector field represents the intracellular fluid velocity, and the scalar color field represents cytosolic pressure. (b) Cell centroid horizontal displacement over time for the three different channel configurations. (c) The inset shows the cell configuration at two different time points as the cellular bleb expands vertically into the crevice of the top wall modeled as a sawtooth function with low frequency and high amplitude.

First, the bleb neck size is varied to produce protrusions of different sizes. The channel gap width is held fixed at 13 *μ*m; the cell is squeezed to 56% of its diameter. With the choice of parameters for bleb expansion and retraction in Table 1, a blebbing cycle corresponds to approximately 7 seconds. The bleb neck size is varied to 4, 8, and 16 *μ*m and the cell undergoes four cycles of bleb expansion and retraction (Fig. 3a). We report the horizontal displacement of the cell centroid in Fig. 3b. As the bleb expands, the cell centroid moves forward gradually, while during bleb retraction, the centroid quickly moves back due to the fast dynamic of cortex re-formation. The cycles of bleb expansion and retraction give rise to cyclic motion of the horizontal displacement of the cell centroid in time. The frequency of these oscillations is the result of the two leading timescales in the problem: the bleb expansion and the cortex reformation timescales. We define the speed of movement to be the slope of the linear fit of the horizontal displacement of the cell centroid and in all three cases, the cell speed is less than 2 *μ*m/min (or less than 10% of the cell diameter). Only the cell with the largest bleb neck size moves forward in the first 30 seconds; when we investigated whether the forward motion is sustained over longer intervals, we found that the cell continues to move forward at the same speed less than 2 *μ*m/min (see Fig. A1 in the Appendix). Based on these results we conclude that cell locomotion is not significant when a cell is confined in a rigid straight wall channel, even with larger forward protrusions.

Next, we assess whether different physical properties of the environment can lead to sustained migration. The width of the channel is varied in Fig. 3c and the corresponding horizontal displacement of the cell centroid is shown in Fig. 3d. As before, the speed is computed as the slope of the linear fit of the horizontal displacement of the cell centroid. Yet again, the cell speed is less than 2 *μ*m/min for all three confinement levels indicating that the cell moves less than 10% of its cell diameter over the course of 4 bleb cycles. Lastly, the geometry of the top channel is altered in order to mimic gaps and pores of the extracellular matrix while the bottom channel is kept straight. This was chosen to resemble the experimental setup in [29]. The top channel is modeled using a rigid sawtooth function of varying frequency (and amplitude) (Fig. 4a). The horizontal displacement of the cell centroid is reported in Fig. 4b. Unlike in the straight channel, we note that the cell can undergo vertical deformations as it expands into the crevices of the top wall and consequently, the centroid displacement is a nonlinear function over time. The largest initial displacement is observed with a low frequency, high amplitude sawtooth channel. By comparison, in the low frequency sawtooth channel, the large gaps in the channel wall allow a bleb to wedge into the gap (inset Fig. 4b). The cell remains in the gap after bleb retraction because the steric interactions with the wall prohibit backward motion. However, we observe that during subsequent bleb cycles, the bleb expands into the gap instead of the channel wall, with no further forward motion after approximately 7 s. This process is the reason the low amplitude oscillations observed in the centroid displacement in the horizontal directions in Fig. 4b (purple line).

### 3.2. Swimming Emerges with Actin Turnover in Blebbing Cells

In the previous section, we probe the cell’s ability to migrate by pressure-driven blebs in confinement. Either changing the protrusion size or the configuration of the physical environment does not produce substantial sustained forward displacement. Introducing additional localized rear contraction to increase the intracellular fluid flow does not greatly improve the results presented here (data not shown). Without introducing cell-surface adhesion, we explore the effect of introducing actin turnover in the thin actomyosin cortex. To model the effect of actin turnover, the constitutive law for the actomyosin cortex needs to be altered to introduce stress relaxation. The cortex is modeled as a thin viscoelastic material with relaxation timescale *λ*, while all other constitutive laws and parameters remain constant. A small strain relaxation timescale is indicative of fast actin turnover and a more viscous, fluid-like response. In the limit of infinite relaxation timescale, an elastic response of the material is recovered. When compared to an elastic cortex, a cell with a viscoelastic cortex deforms irreversibly during cycles of bleb expansion and retraction (Fig. 5). The irreversibility of the motion is due to deformations of the material, as the intracellular fluid flows due to cycles of bleb expansion and contraction, coupled with the evolution of the reference configuration to track the current configuration over time. The faster the relaxation timescale, the more horizontal displacement is observed. In the limit of very slow relaxation (i.e., *λ* → ∞), the elastic response is recovered and the motion is reversible (top panel in Fig. 5). Taken together, these observations suggest that one mechanism to produce adhesion-free motion in confinement is to model the actomyosin cortex as a viscoelastic material.

**Figure 5.**
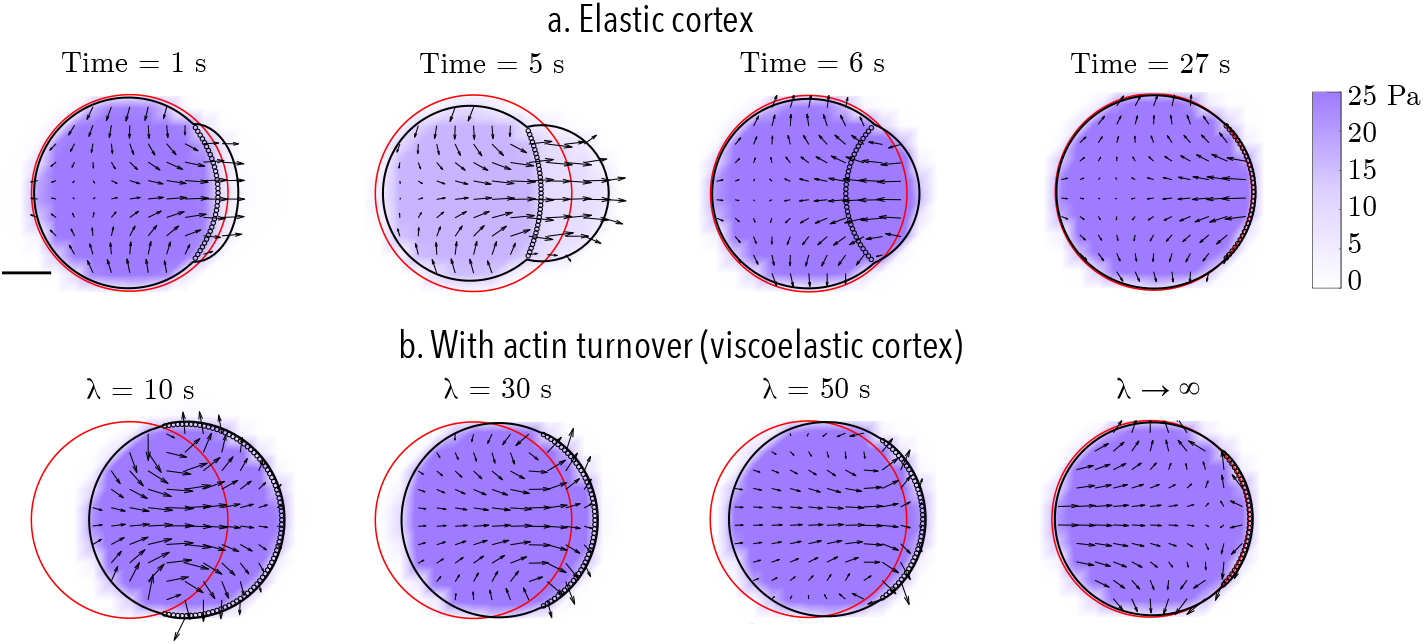
Reciprocity of bleb-based motion for an elastic and a viscoelastic actomyosin cortex with actin turnover timescale *λ*. (a) In the absence of confinement, the motion of bleb expansion and retraction over one cycle (approximately 7 s) and at a time value after 4 bleb cycles (27 s) is (nearly) reciprocal. (b) Cell position after 4 bleb cycles (27 s) for a cell with viscoelastic cortex with different strain relaxation timescales *λ*. Across all panels, the initial bleb neck size is 16 *μ*m. The initial position of the membrane is shown in red. The current position of the membrane is shown in black, and the position of the old cortex are denoted with black circles. The vector field represents the intracellular fluid velocity, and the scalar color field represents cytosolic pressure.

### 3.3. Confinement Enhances Migration Speed of Blebbing Cells

We assess whether confined locomotion is possible with the viscoelastic description of the actomyosin cortex. A blebbing cell with a viscoelastic cortical layer is placed in a confined environment with a rigid straight bottom channel wall and either rigid straight or rigid sawtooth top channel wall (Fig. 6). We report an increase in horizontal speed in all cases over the elastic description of the actomyosin cortex. For the rigid straight channel simulations, the speed is computed through a linear fit of the horizontal displacement over 4 full bleb cycles (27 s). The speed is 2 *μ*m/min for a confinement gap width of 18 *μ*m, 5 *μ*m/min for a width of 13 *μ*m, and 1 *μ*m/min for the narrowest channel width of 8 *μ*m (Fig. 6 a-c respectively). In the case of the irregularly shaped channel, due to the nonlinearity of the centroid horizontal displacement, we report the speed resulting from a linear fit of the horizontal displacement for 60 s rather than 30 s. The speed is 7 *μ*m/min for a high frequency, low amplitude sawtooth, 12 *μ*m/min for a mixed frequency sawtooth, and 9 *μ*m/min (with a stall) for a low frequency, high amplitude sawtooth top channel (Fig. 6 d-f respectively). The trends are similar to the ones in the original model with an elastic cortex. Namely, there is a nonlinear response between confinement gap and migration speed for the rigid straight channel walls. The maximal speed is attained at the intermediate gap level of 13 *μ*m. Another trend is that the cell stalls as it traverses in a channel with a low frequency, high amplitude sawtooth top wall (see Fig. 6f). Similar to simulations with an elastic cortex (data shown in inset Fig. 4c), the bleb expands into a large gap. During subsequent bleb cycles, the cells expands into the gap rather than forward into the channel. No forward motion is observed after approximately 30 s. Interestingly, simulations with an elastic cortex show the cell reaching its steady state location faster than in simulations with a viscoelastic cortex.

**Figure 6.**
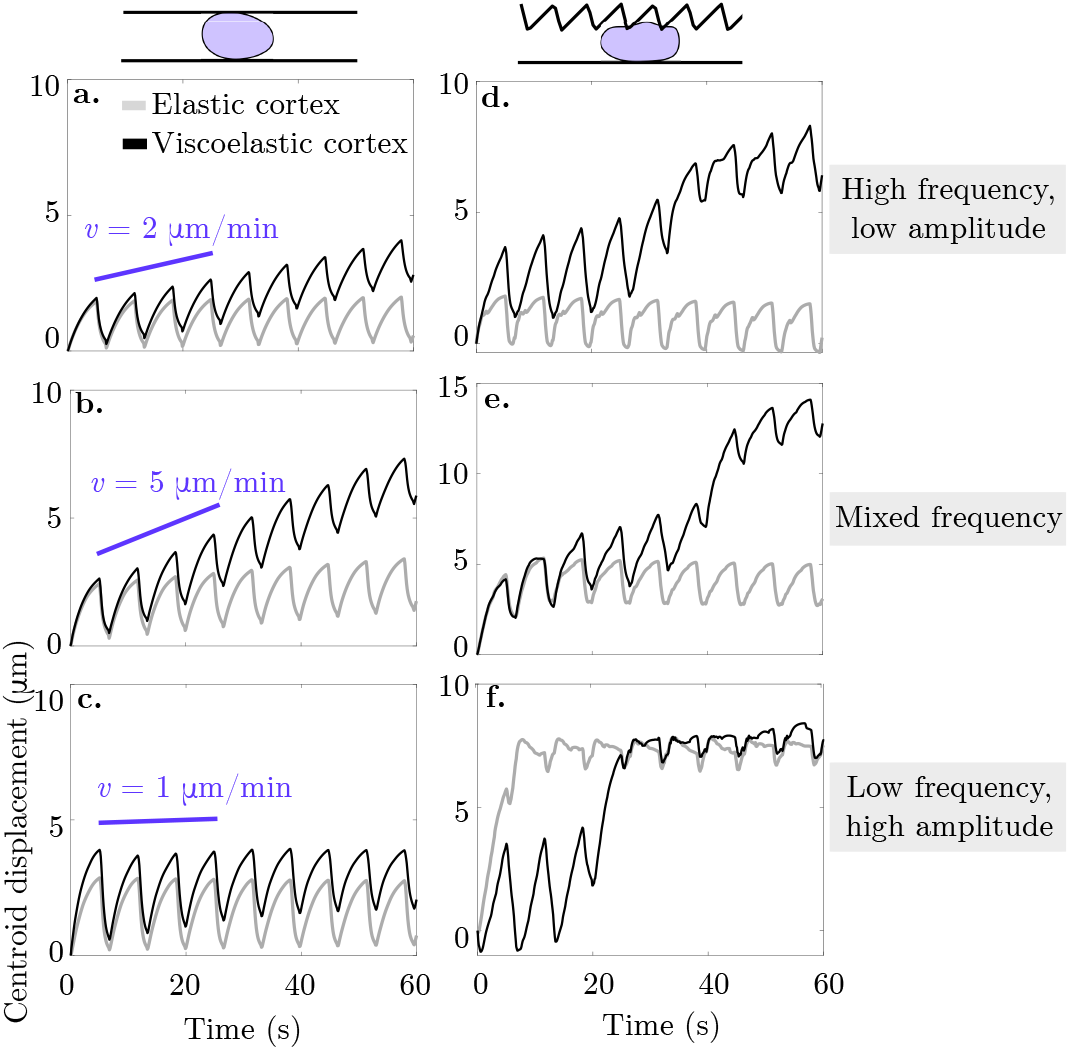
Cortical turnover enables cell motion in a variety of rigid confined environments. The curves represent the horizontal displacement of the cell centroid over time in a confined environment. In panels (a)-(c), the cell in placed in a rigid straight micro-channel of width 18, 13, and 9 *μ*m respectively. Panels (d)-(f) show the horizontal displacement of the cell centroid over time for a cell placed in a asymmetric rigid channel with a straight bottom wall and a: (d) high frequency, low amplitude, (e) mixed frequency, and (f) low frequency, high amplitude sawtooth top wall. Across all panels, bleb neck size is 16 *μ*m. The light grey curve represents the data for an elastic membrane and cortex, while the black curve denotes a viscoelastic cortex with strain relaxation timescale *λ =* 30 s.

Results summarizing the migration speed in various environments as a function of the strain relaxation timescale of the viscoelastic blebbing cell are shown in Fig. 7. A small strain relaxation timescale is indicative of fast actin turnover and a more viscous, fluid-like response. In the limit of infinite relaxation timescale, an elastic response of the material is recovered. Three physical environments are considered: cell placed in free unconfined space, a rigid straight channel with gap width of 13 *μ*m, and a rigid channel with a straight bottom wall and a high frequency, low amplitude sawtooth top wall. In both free space and a rigid straight channel, the cell speed increases monotonically with faster turnover in the actin cortex. For the geometrically asymmetric channel, the cell speed also increases with faster turnover, but there is a sudden jump in speed around *λ =* 30 s. We find that for fast actin turnover, *λ* < 30 s, the cell moves fastest in an asymmetrical rigid channel, while for slower turnover the cell moves fastest in a straight rigid channel. Overall, the trend in our model is that a blebbing cell in a confined environment moves faster than a blebbing cell in free space.

**Figure 7.**
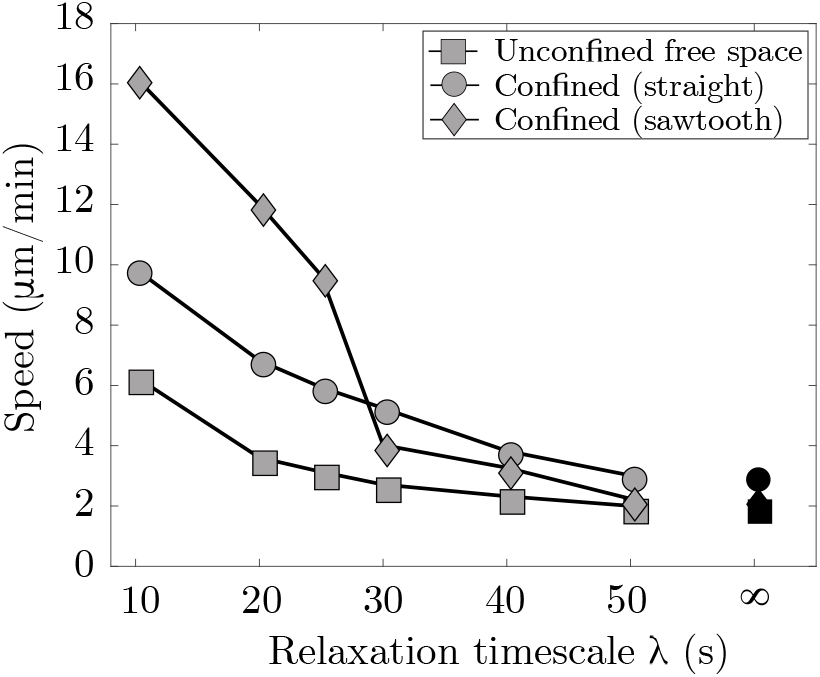
Confinement enhances locomotion speed. Cell speed as a function of strain relaxation timescale for actin turnover, *λ*. Three physical environments are considered: cell placed in free space (unconfined), a straight rigid channel with channel width of 13 *μ*m (confined), and an asymmetric rigid channel with a low amplitude, high frequency sawtooth top channel and a straight bottom channel. Each data point represents the speed computed from a linear fit over 4 full cycles of the horizontal displacement of the cell centroid. The result for no actin turnover (i.e. an elastic cortex) is for *λ →* ∞.

## 4. Discussion and Conclusions

Model simulations of a blebbing cell with a poroelastic cortex immersed in viscous fluid (and no channel) show that membrane shape changes appear reciprocal during bleb expansion and retraction. As a result, a cell cannot swim in zero Reynolds number flow [36]. Even when a geometric asymmetry is introduced with a curved top channel wall, the cell cannot efficiently migrate. In this case, some motion is possible if a bleb wedges the cell into a gap, but sustained motion does not appear possible under our simplified model assumptions. Our results are contradictory to those in [44], where swimming between two straight walls with an elastic cortex was observed. One notable difference between our model and [44] is the force balance on the cortex (Eq. (7)). The cortex model from [44] was studied in [53], where the authors found an imbalance in cortical forces led to large pressure relief compared to the model from [50] that forms the basis for this work. In [32], the authors found that bleb-based migration without substrate adhesion was possible in a discontinuous environment. We did not consider this type of environment here, but migration in our model may be possible when the cortex is treated as an elastic material and both walls are replaced with spaced point sources.

Our results show that the combination of blebbing with cortical actin turnover results in swimming in the absence of confinement. In the context of viscoelasticity, the strain on the cortex evolves in time thus, it creates an asymmetry in the resistance of the material during the bleb expansion and retraction phases. In the case of straight channel walls, we found the fastest migration speed at an intermediate gap size of 13 *μ*m (Fig. 6). In [44], the authors attribute similar results to an increase in intracellular pressure that causes a bleb to form at the rear of the cell. In our model, we only allow blebs to form at the front of the cell and specify that a bleb retracts after 5 s, regardless of the geometry. If we altered this rule to take membrane speed or diffusion of actin into the bleb, it is possible that the cell speed would be affected.

We found that migration speed increases as cortical turnover increases (relaxation timescale *λ* decreases). Interestingly, simulation results show that when *λ* is less than 30 s, cell migration speed is fastest in a geometry when one wall is described by a sawtooth function and the other wall is flat (Fig. 7). As *λ* increases and approaches the case of an elastic cortex, the optimal channel geometry for migration is two flat channel walls. We also observe channel geometries where the cell migrates for a period of time before becoming lodged within a gap, even when the cortex is modeled as a viscoelastic material (see Fig. 6, bottom right). These results point to a non-trivial relationship between the rheology of the cell and its environment. The notion that surface flows can drive adhesion independent migration is not novel to our work [29]; the authors in [54] found that surface treadmilling controlled by active RhoA at the cell rear is sufficient to drive directional cellular motility on 2D surfaces and in liquid.

Our model has several limitations that we hope to address in the future. In order to simulate a viscoelastic material, we re-mesh the membrane and cortex structures after each bleb cycle. The reference configuration can become under-resolved in areas, leading to errors in force computations. In particular, when cortical turnover is fast, the cortex begins to transition from a solid to a fluid, and our algorithm breaks down. In this case, it may be appropriate to simulate the cortex as a fluid. A future direction is to develop better numerical methods to re-mesh the deforming, moving structures and to simulate a thin viscous fluid film immersed in another fluid (such as in [55]).

A cycle of adhesion-independent bleb-based amoeboid motility is thought to consist of protrusion (bleb), outward forces against the channel or extracellular matrix, and followed by a spatially localized rear contraction [56]. We have conducted limited studies to explore the effect of introducing localized myosin-driven contraction, but the focus of this manuscript is on the locomotion due to the pressure gradients induced by blebbing. Rear contraction leads to cortical flows that exacerbate the numerical issues previously mentioned. We hope to comment in the future about the effect of tangential and bulk rear local contractions and the timing of contractions in relation to bleb expansion.

In the present work, the channel is modeled as a rigid structure. In several experiments of confined adhesion-independent migration, one channel wall is a glass coverslip, and the other channel wall is agarose gel [29], which has some elastic properties. It has been suggested that pushing and deforming the channel walls aids the cell’s ability to migrate [24]. One possible extension of this work is to explore how the material properties of the external environment affect cell migration without specific adhesion.

Finally, a limitation of our model is that it is 2D. The relative simplicity of a 2D model allows us to perform simulations over a range of parameter values. Although impressive 3D simulations of amoeboid cell motility were performed in [57,58], they do not include bleb-based motility. A long-term goal is to develop a computationally feasible 3D model that could be used for quantitative comparison to experimental data.

## Author Contributions

Both authors contributed equally in to the conceptualization, methodology, software development, validation, formal analysis of the research. Both authors contributed equally to writing the paper. Both authors have read and agreed to the published version of the manuscript.

## Acknowledgments

This work made use of the High Performance Computing Resource in the Core Facility for Advanced Research Computing at Case Western Reserve University. The authors thank Alex Mogilner (New York University) and Robert Guy (University of California Davis) for useful comments and discussions.

---

**Figure A1.**
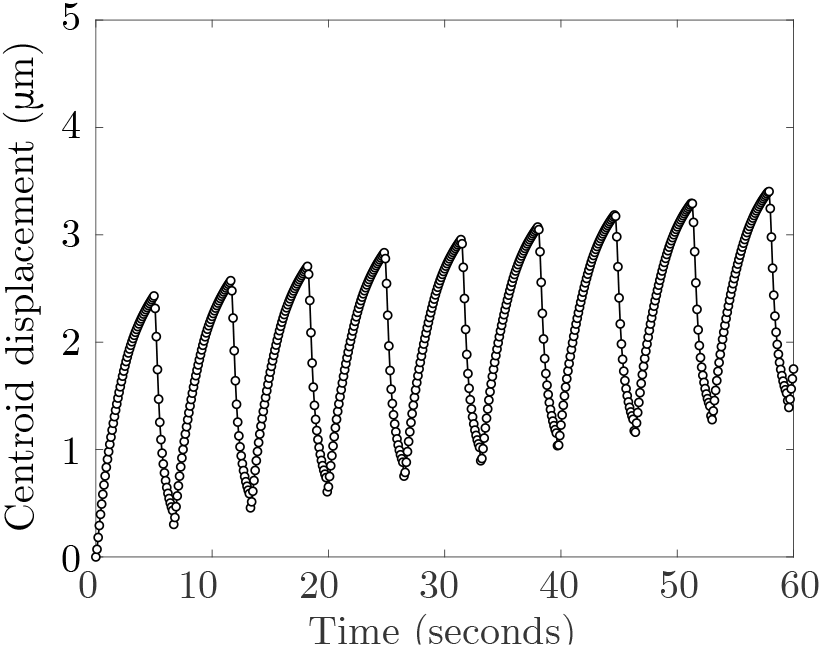
The centroid displacement in the horizontal direction over 60 seconds of a cell with a bleb neck size of 16 *μ*m confined in a straight rigid channel with gap width of 13 *μ*m.

## References

1. Friedl, P.; Wolf, K. Plasticity of cell migration: a multiscale tuning model. J. Cell Biol. 2010, 188, 11–19. doi:10.1083/jcb.200909003.

2. Petrie, R.J.; Doyle, A.D.; Yamada, K.M. Random versus directionally persistent cell migration. Nat. Rev. Mol. Cell Biol. 2009, 10, 538–549. doi:10.1038/nrm2729.

3. Kaye, G.; Siegel, L.; Pascal, R. Cell replication of mesenchymal elements in adult tissues. I. The replication and migration of mesenchymal cells in the adult rabbit dermis. Anat. Rec. 1971, 169, 593–611. doi:10.1002/ar.1091690309.

4. Maaser, K.; Wolf, K.; Klein, C.E.; Niggemann, B.; Zänker, K.S.; Bröcker, E.B.; Friedl, P. Functional hierarchy of simultaneously expressed adhesion receptors: integrin *α*2*β*1 but not CD44 mediates MV3 melanoma cell migration and matrix reorganization within three-dimensional hyaluronan-containing collagen matrices. Mol. Biol. Cell 1999, 10, 3067–3079. doi:10.1091/mbc.10.10.3067.

5. Grinnell, F. Fibroblast mechanics in three-dimensional collagen matrices. J. Bodyw. Mov. Ther. 2008, 12, 191–193. doi:10.1016/j.jbmt.2008.03.005.

6. Friedl, P.; Borgmann, S.; Bröcker, E.B. Amoeboid leukocyte crawling through extracellular matrix: lessons from the *Dictyostelium* paradigm of cell movement. J. Leukoc. Biol 2001, 70, 491–509. doi:10.1189/jlb.70.4.491.

7. Lämmermann, T.; Sixt, M. Mechanical modes of ‘amoeboid’ cell migration. Curr. Op. Cell Biol. 2009, 21, 636–644. doi:10.1016/j.ceb.2009.05.003.

8. Lämmermann, T.; Germain, R.N. The multiple faces of leukocyte interstitial migration. Semin. Immunopathol. 2014, 36, 227–251. doi:10.1007/s00281-014-0418-8.

9. Richardson, B.E.; Ruth, L. Mechanisms guiding primordial germ cell migration: strategies from different organisms. Nat. Rev. Mol. Cell Biol. 2010, 11, 37–49. doi:10.1038/nrm2815.

10. Ibo, M.; Srivastava, V.; Robinson, D.N.; Gagnon, Z.R. Cell blebbing in confined microfluidic environments. PLoS One 2016, 11, e0163866. doi:10.1371/journal.pone.0163866.

11. Srivastava, N.; Kay, R.R.; Kabla, A.J. Method to study cell migration under uniaxial compression. Mol. Biol. Cell 2017, 28, 809–816. doi:10.1091/mbc.E16-08-0575.

12. Gong, X.; Didan, Y.; Lock, J.G.; Strömblad, S. KIF13A-regulated RhoB plasma membrane localization governs membrane blebbing and blebby amoeboid cell migration. EMBO J 2018, 37. doi:10.15252/embj.201898994.

13. Paluch, E.K.; Aspalter, I.M.; Sixt, M. Focal adhesion–independent cell migration. Annu. Rev. Cell Dev. Biol. 2016, 32, 469–490. PMID: 27501447, doi:10.1146/annurev-cellbio-111315-125341.

14. Liu, Y.J.; Le Berre, M.; Lautenschlaeger, F.; Maiuri, P.; Callan-Jones, A.; Heuzé, M.; Takaki, T.; Voituriez, R.; Piel, M. Confinement and low adhesion induce fast amoeboid migration of slow mesenchymal cells. Cell 2015, 160, 659–672. doi:10.1016/j.cell.2015.01.007.

15. Lämmermann, T.; Bader, B.L.; Monkley, S.J.; Worbs, T.; Wedlich-Söldner, R.; Hirsch, K.; Keller, M.; Förster, R.; Critchley, D.R.; Fässler, R.; Sixt, M. Rapid leukocyte migration by integrin-independent flowing and squeezing. Nature 2008, 453, 51–55. doi:10.1038/nature06887.

16. Diz-Muñoz, A.; Krieg, M.; Bergert, M.; Ibarlucea-Benitez, I.; Muller, D.J.; Paluch, E.; Heisenberg, C.P. Control of directed cell migration *in vivo* by membrane-to-cortex attachment. PLoS Biol. 2010, 8, e1000544. doi:10.1371/journal.pbio.1000544.

17. Sahai, E.; Marshall, C.J. Differing modes of tumour cell invasion have distinct requirements for Rho/ROCK signalling and extracellular proteolysis. Nat. Cell Biol. 2003, 5, 711–719. doi:10.1038/ncb1019.

18. Moreau, H.D.; Piel, M.; Voituriez, R.; Lennon-Duménil, A.M. Integrating Physical and Molecular Insights on Immune Cell Migration. Trends Immunol. 2018, 39, 632–643. doi:10.1016/j.it.2018.04.007.

19. Reversat, A.; Gaertner, F.; Merrin, J.; Stopp, J.; Tasciyan, S.; Aguilera, J.; de Vries, I.; Hauschild, R.; Hons, M.; Piel, M.; Callan-Jones, A.; Voituriez, R.; Sixt, M. Cellular locomotion using environmental topography. Nature 2020, 582, 582–585. doi:10.1038/s41586-020-2283-z.

20. Renkawitz, J.; Kopf, A.; Stopp, J.; de Vries, I.; Driscoll, M.K.; Merrin, J.; Hauschild, R.; Welf, E.S.; Danuser, G.; Fiolka, R.; Sixt, M. Nuclear positioning facilitates amoeboid migration along the path of least resistance. Nature 2019, 568, 546–550. doi:10.1038/s41586-019-1087-5.

21. Pflicke, H.; Sixt, M. Preformed portals facilitate dendritic cell entry into afferent lymphatic vessels. J. Exp. Med. 2009, 206, 2925–2935. doi:10.1084/jem.20091739.

22. Thiam, H.R.; Vargas, P.; Carpi, N.; Crespo, C.L.; Raab, M.; Terriac, E.; King, M.C.; Jacobelli, J.; Alberts, A.S.; Stradal, T.; Lennon-Dumenil, A.M.; Piel, M. Perinuclear Arp2/3-driven actin polymerization enables nuclear deformation to facilitate cell migration through complex environments. Nat. Comm. 2016, 7, 10997. doi:10.1038/ncomms10997.

23. van den Berg, M.C.W.; MacCarthy-Morrogh, L.; Carter, D.; Morris, J.; Ribeiro Bravo, I.; Feng, Y.; Martin, P. Proteolytic and opportunistic breaching of the basement membrane zone by immune cells during tumor initiation. Cell Rep. 2019, 27, 2837–2846.e4. doi:10.1016/j.celrep.2019.05.029.

24. Paluch, E.K.; Raz, E. The role and regulation of blebs in cell migration. Curr. Op. Cell Biol. 2013, 25, 582–590. doi:10.1016/j.ceb.2013.05.005.

25. Malawista, S.E.; de Boisfleury Chevance, A.; Boxer, L.A. Random locomotion and chemotaxis of human blood polymorphonuclear leukocytes from a patient with leukocyte adhesion deficiency-1: normal displacement in close quarters via chimneying. Cell Mot. 2000, 46, 183–189. doi:10.1002/1097-0169(200007)46:3<183::AID-CM3>3.0.CO;2-2.

26. Charras, G.T.; Coughlin, M.; Mitchison, T.J.; Mahadevan, L. Life and times of a cellular bleb. Biophys. J. 2008, 94, 1836–1853. doi:10.1529/biophysj.107.113605.

27. Charras, G.T.; Yarrow, J.C.; Horton, M.A.; Mahadevan, L.; Mitchison, T.J. Non-equilibration of hydrostatic pressure in blebbing cells. Nature 2005, 435, 365–369. doi:10.1038/nature03550.

28. Tinevez, J.Y.; Schulze, U.; Salbreux, G.; Roensch, J.; Joanny, J.F.; Paluch, E. Role of cortical tension in bleb growth. Proc. Natl. Acad. Sci. USA 2009, 106, 18581–18586. doi:10.1073/pnas.0903353106.

29. Bergert, M.; Erzberger, A.; Desai, R.A.; Aspalter, I.M.; Oates, A.C.; Charras, G.; Salbreux, G.; Paluch, E.K. Force transmission during adhesion-independent migration. Nat. Cell Biol. 2015, 17, 524–529. doi:10.1038/ncb3134.

30. Maugis, B.; Brugués, J.; Nassoy, P.; Guillen, N.; Sens, P.; Amblard, F. Dynamic instability of the intracellular pressure drives bleb-based motility. J. Cell Sci. 2010, 123, 3884–3892. doi:10.1242/jcs.065672.

31. Maxian, O.; Mogilner, A.; Strychalski, W. Computational estimates of mechanical constraints on cell migration through the extracellular matrix. PLoS Comput. Biol. 2020, 16, e1008160. doi:10.1371/journal.pcbi.1008160.

32. Tozluoğlu, M.; Tournier, A.L.; Jenkins, R.P.; Hooper, S.; Bates, P.A.; Sahai, E. Matrix geometry determines optimal cancer cell migration strategy and modulates response to interventions. Nat. Cell Biol. 2013, 15, 751–762. doi:10.1038/ncb2775.

33. Zhu, J.; Mogilner, A. Comparison of cell migration mechanical strategies in three-dimensional matrices: a computational study. Int. Focus 2016, 6, 20160040. doi:10.1098/rsfs.2016.0040.

34. Prost, J.; Jülicher, F.; Joanny, J.F. Active gel physics. Nat. Phys. 2015, 11, 111–117. doi:10.1038/NPHYS3224.

35. Hawkins, R.; Piel, M.; Faure-Andre, G.; Lennon-Dumenil, A.; Joanny, J.; Prost, J.; Voituriez, R. Pushing off the walls: a mechanism of cell motility in confinement. Phys. Rev. Lett. 2009, 11, 058103. doi:10.1103/PhysRevLett.102.058103.

36. Purcell, E. Life at low Reynolds number. Am. J. Phys. 1977, 45, 3–11. doi:10.1119/1.10903.

37. Brennen, C.; Winet, H. Fluid mechanics of propulsion by cilia and flagella. Annu. Rev. Fl. Mech. 1977, 9, 339–398. doi:10.1146/annurev.fl.09.010177.002011.

38. Gray, J.; Hancock, G.J. The propulsion of sea-urchin spermatozoa. J. Exp. Biol. 1955, 32, 802–814. doi:10.1242/jeb.32.4.802.

39. Barry, N.; Bretscher, M. *Dictyostelium* amoebae and neutrophils can swim. Proc. Natl. Acad. Sci. U.S.A. 2010, 107, 11376–11380. doi:10.1073/pnas.1006327107.

40. Othmer, H. Eukaryotic cell dynamics from crawlers to swimmers. WIREs Comput. Mol. Sci. 2019, 9, e1376. doi:10.1002/wcms.1376.

41. Wang, Q.; Othmer, H. Analysis of a model microswimmer with applications to blebbing cells and mini-robots. J. Math. Biol. 2018, 76, 1699–1763. doi:10.1007/s00285-018-1225-y.

42. Wang, Q.; Othmer, H. Computational analysis of amoeboid swimming at low Reynolds number. J. Math. Biol. 2016, 72, 1893–1926. doi:10.1007/s00285-015-0925-9.

43. Wu, H.; de León, M.A.P.; Othmer, H.G. Getting in shape and swimming: the role of cortical forces and membrane heterogeneity in eukaryotic cells. J. Math Biol 2018, 77, 595–626. doi:10.1007/s00285-018-1223-0.

44. Lim, F.; Koon, Y.; Chiam, K.H. A computational model of amoeboid cell migration. Comp. Meth. Biomech. Biomed. Eng. 2013, 16, 1085–1095. PMID: 23342988, doi:10.1080/10255842.2012.757598.

45. Cortez, R. The method of regularized Stokeslets. SIAM J. Sci. Comput. 2001, 23, 1204–1225. doi:10.1137/S106482750038146X.

46. Yip, A.K.; Chiam, K.H.; Matsudaira, P. Traction stress analysis and modeling reveal that amoeboid migration in confined spaces is accompanied by expansive forces and requires the structural integrity of the membrane–cortex interactions. Int. Biol. 2015, 7, 1196–1211. doi:10.1039/c4ib00245h.

47. Salbreux, G.; Charras, G.; Paluch, E. Actin cortex mechanics and cellular morphogenesis. Trends Cell Biol. 2012, 22, 536–545. doi:10.1016/j.tcb.2012.07.001.

48. Gardel, M.; Kasza, K.; Brangwyne, C.; Liu, J.; Weitz, D. Mechanical response of cytoskeletal networks. Met. Cell Biol. 2008, 89, 487–519. doi:10.1016/S0091-679X(08)00619-5.

49. Copos, C.; Guy, R. A porous viscoelastic model for the cell cytoskeleton. ANZIAM J 2018, 59, 472–498. doi:10.1017/S1446181118000081.

50. Strychalski, W.; Guy, R.D. A computational model of bleb formation. Math. Med. Biol. 2013, 30, 115–130. doi:10.1093/imammb/dqr030.

51. Moeendarbary, E.; Valon, L.; Fritzsche, M.; Harris, A.R.; Moulding, D.A.; Thrasher, A.J.; Stride, E.; Mahadevan, L.; Charras, G.T. The cytoplasm of living cells behaves as a poroelastic material. Nat. Mater. 2013, 12, 253–261. doi:10.1038/nmat3517.

52. Strychalski, W.; Guy, R.D. A computational model of bleb formation. Math. Med. Biol. 2013, 30, 115–130. doi:10.1093/imammb/dqr030.

53. Maxian, O.; Strychalski, W. 2D force constraints in the method of regularized Stokeslets. Comm. App. Math. Comp. Sci. 2019, 14, 149–174. doi:10.2140/camcos.2019.14.149.

54. O’Neill, P.R.; Castillo-Badillo, J.A.; Meshik, X.; Kalyanaraman, V.; Melgarejo, K.; Gautam, N. Membrane flow drives an adhesion-independent amoeboid cell migration mode. Dev. Cell 2018, 46, 9–22.e4. doi:10.1016/j.devcel.2018.05.029.

55. Whitfield, C.A.; Hawkins, R.J. Immersed boundary simulations of active fluid droplets. PLoS One 2016, 11, e0162474. doi:10.1371/journal.pone.0162474.

56. Charras, G.; Paluch, E. Blebs lead the way: how to migrate without lamellipodia. Nat. Rev. Mol. Cell Biol. 2008, 9, 730–736. doi:10.1038/nrm2453.

57. Campbell, E.J.; Bagchi, P. A computational model of amoeboid cell motility in the presence of obstacles. Soft Matter 2018, 14, 5741–5763. doi:10.1039/c8sm00457a.

58. Moure, A.; Gomez, H. Three-dimensional simulation of obstacle-mediated chemotaxis. Biomech. Model. Mechanobiol. 2018, 17, 1243–1268. doi:10.1007/s10237-018-1023-x.

